# Model to link cell shape and polarity with organogenesis

**DOI:** 10.1101/699413

**Authors:** Bjarke Frost Nielsen, Silas Boye Nissen, Kim Sneppen, Ala Trusina, Joachim Mathiesen

**Affiliations:** Niels Bohr Institute, University of Copenhagen, Blegdamsvej 17, 2100 Copenhagen, Denmark

## Abstract

How do tubes — gut or neural tube — form from flat sheets of polarized cells? The prevalent view is that it is a two-step process: first cells wedge to bend the sheet, then cells intercalate and extend the initial invagination into a tube. We computationally challenged this model by asking if one mechanism (either cell wedging or intercalation) may suffice for the entire sheet-to-tube transition. Using a physical model with epithelial cells represented by polarized point particles, we show that either cell intercalation or wedging alone can be sufficient and each can both bend the sheet and extend the tube. When working in parallel, the two mechanisms increase the robustness of the tube formation. The successful simulations of Drosophila salivary gland, Sea urchin gastrulation and mammalian neurulation support the generality of our results.

## Introduction

Early tubes in embryonic development – gut and neural tubes – form out of epithelial sheets. In mammalian embryos and *Drosophila*, the cell sheet wraps around the tube axis until the edges make contact and fuse. As a result of such *wrapping*, a tube is formed parallel to the sheet. In sea urchin, the gut is formed orthogonal to the epithelial plane by *budding* out of the plane. Budding also appears to be a predominant form of tube formation in organ development (lungs, kidneys, salivary gland and trachea in *Drosophila* [1]. Both wrapping and budding sheet-to-tube transitions are driven by the same key mechanisms: changes in cell shape, e.g. cell wedging by changes in apical relative to the basal surfaces – apical constriction (AC) [2] or basal constriction [3, 4]; contracting myosin cables spanning across cells and Convergent Extension (CE) by directed cell intercalation [1, 5]. (In the following, we will refer to apical or basal constriction as *wedging* and directed cell intercalation as CE).

Until recently, the consensus has been that wedging and CE each lead to distinct morphological transformations: wedging bends the sheet and is a primary mechanism for invagination in budding [6] and CE elongates the sheet and the eventual tube [1]. However, results by Chung et al. [5] and Sanchez-Corrales et al. [7] show that invagination in *Drosophila* salivary gland can happen in the absence of wedging and may result from radial CE. Furthermore, Nishimura et al. [8] suggested that In mammalian neurulation CE and wedging are coupled. They argue that Planar Cell Polarity (PCP) may be mediating this coupling: First, the direction of cell intercalations, orthogonal to the tube axis, is set by PCP. Second, wedging has to be anisotropic – have preferred direction parallel to PCP and intercalations – for the sheet to wrap into a tube and not a sphere. The anisotropy of wedging, however, is rarely considered [8–10], possibly because the reported results are often limited to 2D cross-sectional views. The anisotropy may stem from the coupling between PCP and wedging – both apical *and* basal. This is supported by data at the molecular level (for neural tube closure [8, 11], midbrain-hindbrain boundary in zebrafish [3, 4], gastrulation in *C. elegans* [12], sea urchin [13], and *Xenopus* [14]).

This recent development opens for new questions: *What are wedging and CE capable of on their own? Can invagination by CE happen in systems other than salivary glands? Is anisotropy in wedging important for tubulogenesis and, if so, when?*

We here introduce a theoretical model to address these questions. Theoretical models have been important for understanding tubulogenesis, however they are often limited to 2D and thus focus on either wedging or CE [15–17]. While there are 3D models for budding and neurulation [18, 19], they lack the coupling between PCP, wedging and CE and do not capture the entire sheet-to-tube transition. To close this gap we introduce a model of polarized cell–cell interactions where cells are treated as point particles. As a starting point for our model, we consider the model suggested in [20] which was used to study directional adhesion mediated by apical-basal (AB) polarity and PCP. The model in [20], however, could not explicitly account for changes in cell shapes. Here, we show that the effect of cell wedging can be modeled within a point-particle representation by modifying cell-cell forces to favor a tilt in AB polarities.

In line with data in Chung et al. [5], simulations show that although CE alone can lead to a budding transition, it is less reliable, with frequent failure of invagination and misorientation. Our results suggest that the isotropic wedging orients the budding and allows for robust invagination. When applied to wrapping in neurulation, we find that anisotropic wedging alone was insufficient for final tube closure. However, both closure and tube separation from the epithelium can be aided by differential proliferation. Furthermore, we find that anisotropic wedging on its own may be sufficient for tube elongation. Together, our results support the mutual complementarity of wedging and CE in bending and elongation.

## Results

To investigate the role of cell wedging in budding of salivary gland placoids and neurulation we aimed at capturing both isotropic and anisotropic (PCP-driven) wedging with as few parameters as possible.

### Modelling wedging of a point particle by favoring tilt in AB

Apical constriction leads to cell wedging and as a consequence the AB axes of neighboring cells become tilted towards the wedged cell (Fig 1B–C). In [20] a flat epithelial sheet was modelled by a cell-cell interaction force favouring parallel AB polarities in neighboring cells (Fig 1A, Eq 4). To model the effect of wedging we modify the force to favor AB polarity vectors **p**_*i*_ in neighbor cells to tilt towards the wedged cell (Fig 1B and 1C). That is, when the force is calculated, we replace **p**_*i*_ by 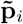 (Eqs 1–3).

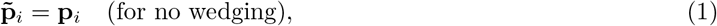

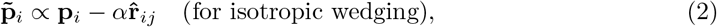

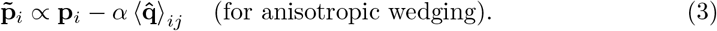

**Fig 1.**
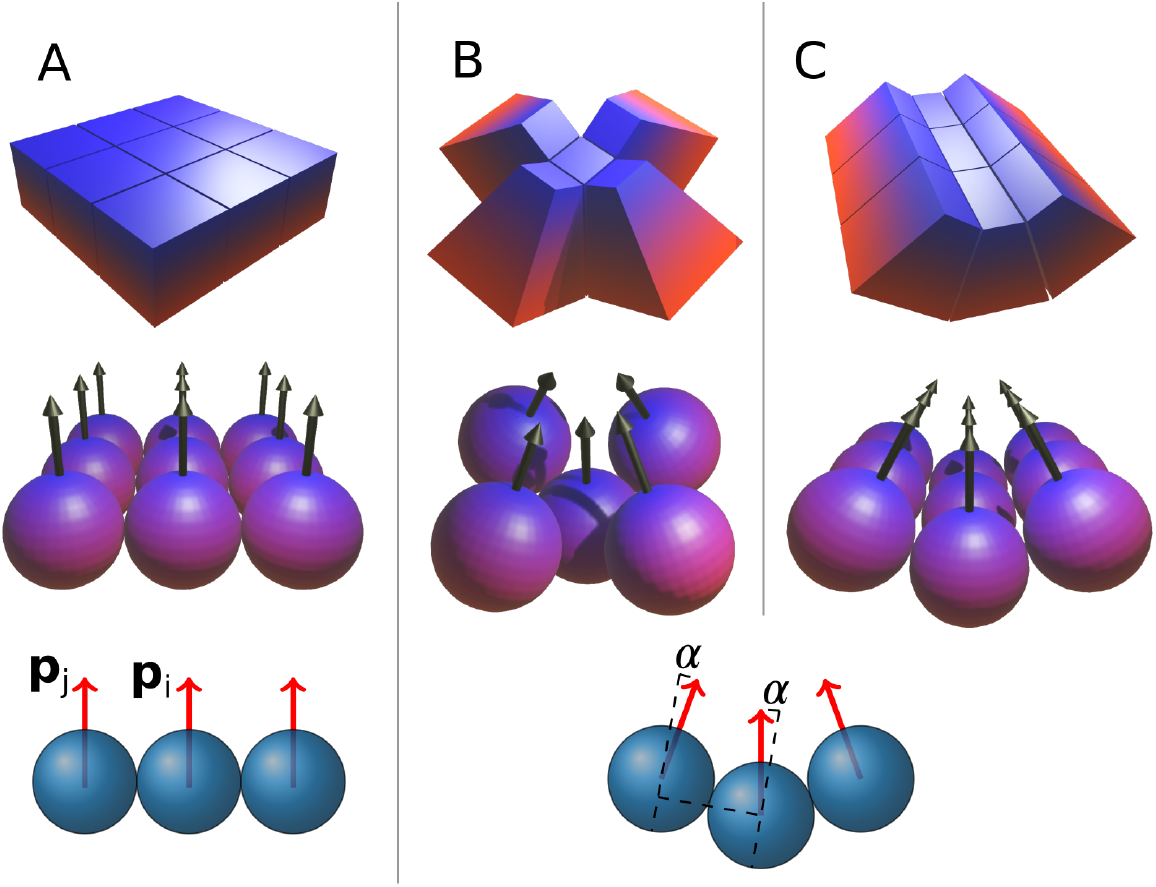
Wedging is introduced through a cell-cell interaction that favors tilted AB polarity vectors. *α* is the extent of the wedging. The blue-red gradient indicates the apical-basal axis. **A**) No wedging (*α* = 0), AB polarities (arrows) tend to be parallel. (**B**) With isotropic wedging, the tilt *α* is the same in all directions. (**C**) With anisotropic wedging, the tilt has a preferred direction. Blue and red signify respectively basal and apical surfaces. *p*_*i*_ and *p*_*j*_ are the AB polarities of cell *i* and *j*.

Here, 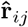 is the normalized displacement vector between cells *i* and *j* while 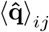 is the average PCP vector of the two interacting particles.

This change required only one parameter, *α*, setting the extent of the tilt (large *α* corresponds to pronounced wedging). If the wedging is isotropic, i.e. equally pronounced in all directions [7], all neighbors to the wedged cell tend to tilt equally. In neurulation, the wedging is anisotropic: the wedging happens primarily parallel to the cell’s PCP and perpendicular to the axis of the tube [8]. To capture this PCP-directed anisotropy, we couple the direction of AB tilting to the orientation of the cell’s PCP (Eq 3, Fig 1C). See the Methods section for details of the model and simulations.

Note, that we aim to only capture the effect of wedging-PCP coupling and not the molecular mechanism. Also, in an attempt to generalize our results, we focus on a minimal set of conditions necessary for the final outcome.

To test the validity of our approach, we first consider the complementary roles of CE and wedging in budding.

### Complementary and unique roles of CE and wedging in budding

Reflecting the viewpoint that tube budding is a two stage process consisting of wedging-driven invagination and subsequent convergent extension, computational models have generally focused on either of the two stages [15–17]. However, to date no computational models have managed to recapitulate the results by [7] and [5] suggesting that CE contributes to invagination and may even drive it in the absence of wedging.

To validate our approach, we set out to reproduce these experimental observations. We start with a flat sheet of AB polarized cells. Motivated by the possible link between organizing signals (e.g. WNT), PCP and wedging [21, 22], we define a region of “organizing signals” such that the cells within this region exhibit isotropic wedging and PCP. In salivary glands, the apically constricting cells are distributed on a disk around the future center of the tube. With this configuration, we did not find parameters where both CE and wedging could act in parallel to form the tube. A two-step process, wedging followed by CE, was able to produce invagination and tube extension. However, a ring of basally constricting cells (with or without apically constricting disk) remedied this problem and allowed for wedging and CE to act in parallel. Supporting this, the data by [7] suggests that there are basally constricting cells in the outer region of the placoid. Furthermore, basal and apical constriction seem to be induced by the same organizing signal [3] through PCP pathways. Also in sea urchin gastrulation, both types of wedging seem to be at play [23]. For simplicity, we limit our simulations to basal wedging, where basally constricting cells are distributed on a ring (Fig 2A and S5 Fig)

**Fig 2.**
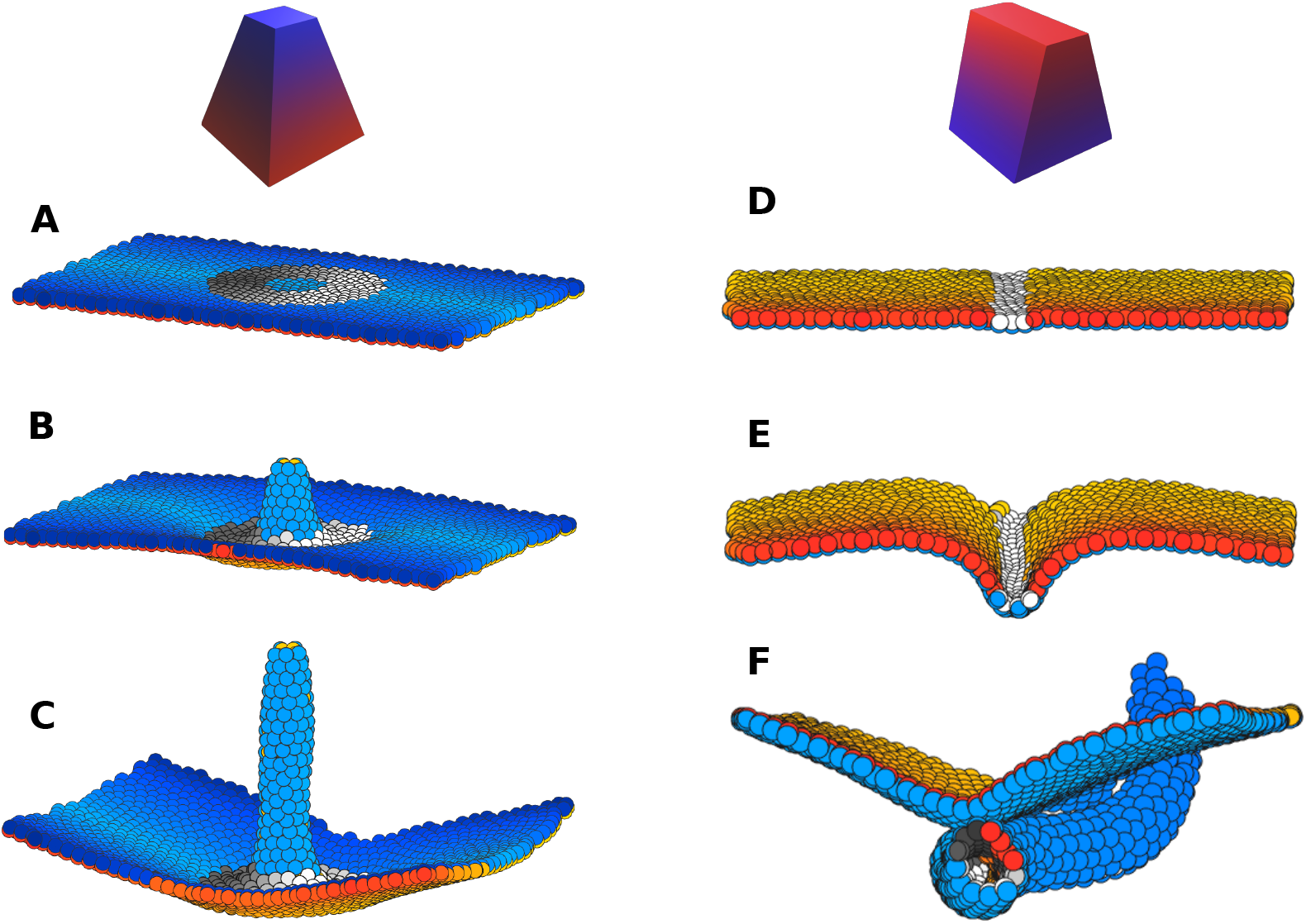
Isotropic and anisotropic wedging drive budding and wrapping, respectively. Wedging cells are labeled in gray, with a shading that indicates the PCP direction. (**A-C**) Time evolution of budding simulation (similar to salivary glands). Here, gray cells constrict basally and all cells on and inside the ring intercalate radially. Here, *λ*_3_ = 0.1, |*α*| = 0.5. (**D-F**) Time evolution of wrapping simulation (similar to neurulation). Here, gray cells representing neuroepithelium, constrict apically and constriction is anisotropic, follows the direction of PCP (Eq 3). Cells proliferate only at the gray/colored boundary (with 7 hour doubling time), mimicking differential proliferation at the neuroepithelium/ectoderm boundary. Here, *λ*_3_ = 0, *|α|* = 0.5. Throughout, *λ*_2_ = 0.4.

Our budding simulations thus show that successful invagination and tube elongation can proceed if both wedging and PCP (and thus CE) act in parallel (S1 Video, Fig 2A-C). We have also succeeded in simulating sea urchin gastrulation where budding starts from a sphere of cells (Fig 3, S5 Video, [24, 25]). This proceeds essentially like in the planar case (see Methods for details).

**Fig 3.**
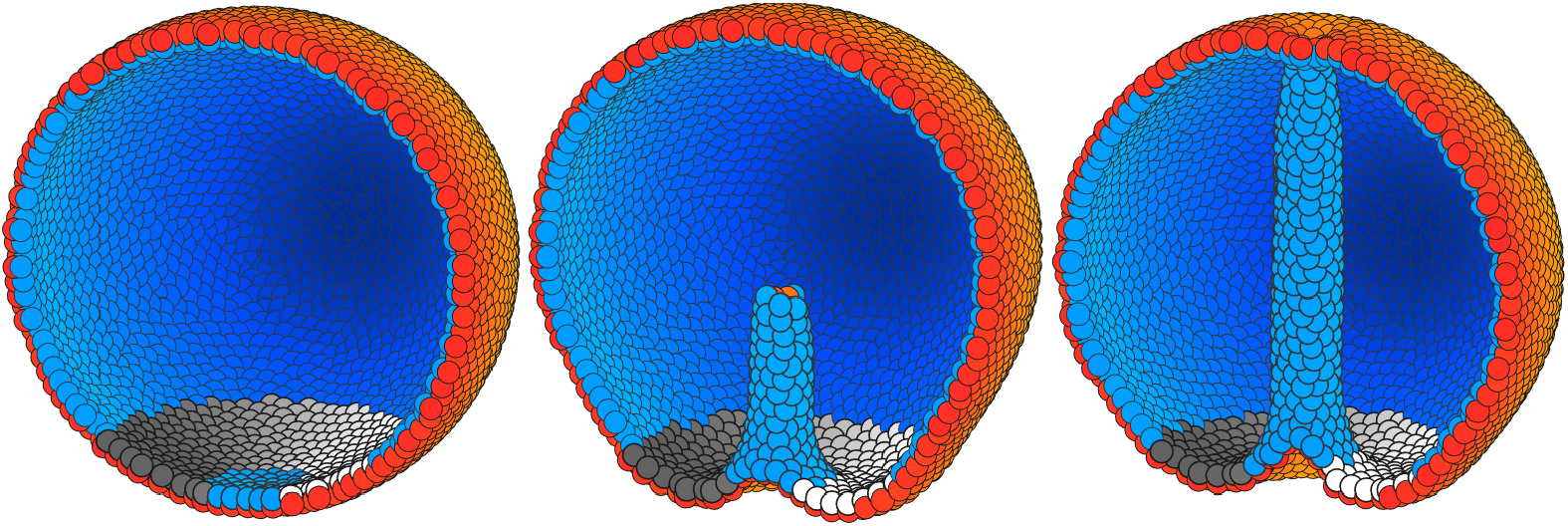
Isotropic wedging in conjunction with PCP is sufficient to drive sea urchin gastrulation without external forcing. The gray ring shows cells with (isotropic) basal constriction and the shading indicates the direction of planar cell polarity, which curls around the vertical axis in our simulation. See section Modeling Gastrulation for details.

In addition, we find that budding *can* proceed without wedging. However, robustness of the outcome decreases in two ways. First, the proportion of failed invaginations is higher (S1 Fig). Second, the tube can form on either side of the epithelial plane.

With loss of apical constriction, noise is necessary to break the symmetry between the two sides of the plane and initiate the CE-driven tubulation in one of the two directions orthogonal to the plane. Thus, it seems that the role of wedging is to aid in the initial invagination and ensure correct orientation. This is a plausible explanation for the results obtained in [5] where the authors knocked-out wedging in the context of *Drosophila* salivary gland formation and observed that the budding process could still proceed, but with reduced reliability and orientational stability. In contrast to our results and the findings by [7], [5] do not consider cell intercalations by CE but propose that supracellular myosin cables drive tissue bending in the absence of wedging. It will be interesting to extend our approach to include an analog of myosin cables through e.g. PCP-coupled supracellular forces, however, it is outside of the scope of the current work.

Cell shape change, intercalation and tissue compression by supracellular myosin cables are also key players in wrapping [8]. The differences that cause some tubes to form parallel and others orthogonal to the epithelial plane appear to be encoded in the geometrical arrangement of the cells that participate in these three processes. In budding such cells are arranged on a ring or a disk (circular symmetry), while in wrapping they are arranged on a band (axial symmetry).

### Anisotropic wedging and differential proliferation are sufficient for wrapping

To test if this difference in geometry alone is sufficient for wrapping, we choose a stripe of cells in the middle of the epithelial sheet to represent the neuroepithelium (NE) (shown by gray in Fig 2D-E) and the remaining cells to represent ectoderm (E) (colored cells in Fig 2D-E). The NE cells are then assigned anisotropic apical constriction and PCP pointing orthogonal to the future tube axis (S4 Fig).

#### Wrapping requires anisotropy in wedging

In the case of isotropic wedging one would expect a collection of NE cells to eventually form a round invagination or spherical lumen — the minimum energy state (S4 Video). If we impose isotropic wedging in our neurulation simulations, a bulging, rounded invagination is observed, see S3 Video.

Motivated by the results of [8], showing that wedging is anisotropic (Eq 3) and cells wedge primarily in the direction orthogonal to the tube axis, we asked if anisotropic wedging can aid in tube closure. As expected, the tissue bends around the tube axis without capping at the ends of the tube (Fig 1C, S2 Fig).

Interestingly, anisotropic wedging also leads to CE cell intercalation, narrowing and elongating neuroepithelium (see S3 Fig), thus supporting the link between PCP-driven wedging and cell intercalations. The simple intuitive argument for this comes from how wedged cells pack in the tube. In the minimum energy state the extent of wedging, *α*, determines how many cells can pack around the tube’s circumference (Fig 1). To minimize energy, the “extra” cells will be displaced along the tube axis (S3 Fig). CE-driven narrowing of the epithelium was proposed to be important for tube closure [26]. In our simulations, wedging and CE alone succeeded in bending the tissue in an axially symmetric fashion (S2 Fig), however, we could not obtain successful tube closure even with maximally possible CE and wedging (both tuned by the strength of *α* in Eq 3). This suggests that additional mechanisms are necessary for final tube closure.

#### Buckling by proliferation at the NE boundary aids in tube closure

Images of neurulation cross-sections (see e.g. [27]) show a strong bending at the neuroepithelium-ectoderm (NE) boundary with the curvature opposite to that inside of neuroepithelium (neural folds) [28]. This is believed to be a result of combined forces from the ectoderm due to i) change in cell shape (ectoderm cells become flatter and neuroepithelial cells become taller); ii) adhesion between basal surfaces of NE and E close to the neuroepithelium-ectoderm (NE-E) boundary [28] and iii) increase in cell density at this boundary either due to cell proliferation or intercalation [29].

Our goal was to test if the model can capture full tube closure with at least one of the mechanisms, so for simplicity, we focused on differential proliferation. When cells were set to proliferate only at the NE-E interface [29], we found that the resulting buckling can lead to successful neural tube closure (S2 Video). In the simulations, the out-of-equilibrium buckling created by rapid cell proliferation is necessary to create a narrow neck that allows epithelial folds to fuse. We find that tubulation is possible within a rather broad range of cell cycles (3h–16h). Shorter *or* longer cell cycles resulted in open-tube morphologies reminiscent of neural tube defects such as spina bifida (Fig 4). In both cases the folds are too far apart to fuse, but for different reasons. If proliferation is too slow, the folds are far apart because the buckling is too weak. On the other hand, when proliferation is too fast, the sheet does not have time to equilibrate and CE does not catch up in narrowing it. Because of this, some sections of the tube become too wide to fuse. Interestingly this can sometimes lead to tube doubling/splitting (S6 Fig).

**Fig 4.**
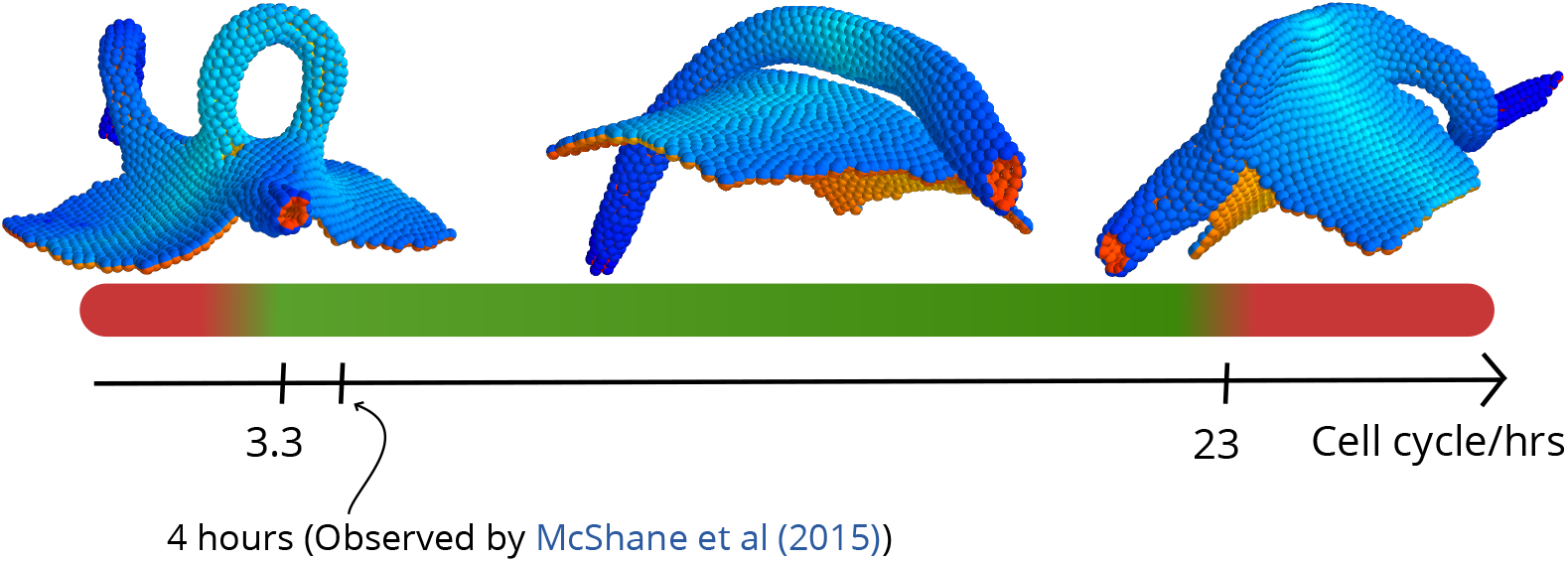
The cell cycle length at the neuroepithelial-ectoderm boundary affects tube closure. For cell cycle lengths below 3.3h and above 23h the neural tube fails to close in our simulations. It should be noted that this broad interval also contains the cell cycle length of 4h found for cells in the DLHP by [29]. The insets show outcomes of simulations run at short (2.6h), intermediate (12h) and long (26h) cell cycle lengths.

The effect of slow proliferation in our simulations is in line with the experimental data. In [30] it was shown that low proliferation rates could lead to neural tube defects in mice. In humans, mutations of the PAX3 transcription factor are implicated in Waardenburg syndrome [31, 32] characterized by incomplete neural tube closure. The same transcription factor has been shown to be essential in ensuring sufficient cell proliferation [33]. The effect of increased (compared to wild-type) proliferation has not been addressed experimentally and we hope that our predictions will motivate experiments in this direction.

## Discussion

Larger organisms rely on tubes for distributing nutrients across the body as well as for exocrine functions. How these tubes reliably form is an open question, but a few recurrent mechanisms are known, e.g. directed or differential proliferation, changes in cell shapes, supracellular myosin cables, directed adhesion and cell rearrangements. As evolution proceeds by tinkering rather than engineering, it is not surprising that these mechanisms have overlapping functions. Recently quantitative experiments [5, 7, 8] enabled us to look beyond a “one mechanism, one function” relationship and towards a map of where mechanisms overlap and how they complement each other.

In this work, we have made a step towards charting the functional overlap and complementarity among CE, wedging, and proliferation. A phenomenological point-particle representation allowed us for the first time to combine PCP-driven cell intercalation (CE) and anisotropic wedging in thousands of cells in 3D and with a few free parameters.

This allowed us to arrive at the following key results. First, our simulations recapitulate that CE can drive invagination in the absence of wedging [5, 34]. Thus suggesting that this is a general mechanism, that it does not require forces from surrounding tissues and that it is also possible in invaginating systems other than the salivary glands in *Drosophila*. The invagination is however unreliable and isotropic wedging plays a complementary role by setting the direction of invagination.

Second, our results predict that anisotropic, PCP-coupled wedging may play a role in tube formation and elongation. Our model predicts that anisotropy in wedging maintains axial symmetry of the tube during wrapping. Remarkably, anisotropic wedging can also effectively result in CE-like cell intercalation and lead to tube elongation. While we have only tested the contribution of anisotropic wedging in wrapping, the same principle may apply in budding, were the reported isotropic wedging [7, 34] seems to become anisotropic along the axis of the tube as it is elongating [35]. It will be interesting to explore this hypothesis theoretically and experimentally. Such isotropic to anisotropic transition in wedging has been reported in *Drosophila* furrow formation [9, 36]. The visual inspection of tube cross sections in the pancreas and kidneys suggest that cells are wedged, and while by analogy to neurulation it is reasonable to expect wedging to be anisotropic, it remains to be confirmed experimentally by e.g. whole mount 3D imaging of stained tubes.

Third, buckling by differential proliferation (faster at the neuroepithelium/ectoderm boundary than in the remaining tissue) together with anisotropic wedging within neuroepithelium is sufficient for tube closure and separation. Differential proliferation has been proposed by [29] as a mechanism for forming DLHP – regions where the tissue curvature has the same sign as at medial hinge points (MHP). We find that modifying the extent of apical constriction or how it is distributed – i.e. throughout entire neuroepithelium, or combinations of DLHPs and MHP, could not result in tube closure. Instead, our results highlight the importance of creating opposite curvature at the boundaries. Our simulations suggest that differential proliferation buckles the boundaries and aids tube closure as it curves the epithelium opposite to the curvature resulting from apical constriction.

Our simulations predict a wide range of proliferation rates capable of producing sufficient buckling for closure. These results call for testing for differential proliferation in systems without DLHP’s (by accelerating or reducing proliferation rate in mutants or by molecular inhibitors [37]). While not immediately feasible, it is also interesting to consider how to perturb the “opposite” curvature by interfering with differences in cell shapes or basal adhesion [28] of the neuroepithelium and ectoderm close to the boundary.

Models of tubulogenesis date back to at least a few decades [38], however most of them are limited to 2D and focus on either wedging or cell intercallation. Recently, Inoue et al. [19] formulated a 3D vertex model of neurulation focusing on cell elongation, apical constriction, and active cell migration. The model does not include either cell proliferation, or PCP, but instead relies on active cell migration to pull the neural cells towards the midline. While successful in bringing folds sufficiently close, it does not cover the separation of the tube from the sheet. Also, a recent 3D model of tube budding in lung epithelium arrived at the conclusion that anisotropic wedging can only result in rounded tubes and is insufficient to drive the entire process [18].

We have demonstrated that cell wedging can be phenomenologically captured in a point-particle representation. This is not restricted to apical constriction but also covers e.g. basal constriction and can, in a similar spirit, be extended to capture changes in cell height and width. This could allow for modeling a wider range of phenomena where morphological changes are driven by these differences. Furthermore, we are now in a position to address tube branching in e.g. lungs and vascularization, where cells forming the tubes also are the ones that secrete organizing signals that locally re-orient PCP polarities and may induce anisotropic changes in cell shapes.

## Methods

### Model

Following [20], cells are treated as point particles interacting with neighboring cells through a pair-potential *V*_*ij*_. The potential has a rotationally symmetric repulsive term and a polarity-dependent attractive term. In terms of *r*_*ij*_ (the distance between two cells *i* and *j*), the dimensionless potential can be formulated as

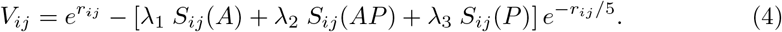

The parameters *λ*_*i*_ are coupling constants which define the strength of polar interactions in the model. *S*_*ij*_(*A*) quantifies the coupling between AB polarity and position, whereas *S*_*ij*_(*AP*) and *S*(*P*)_*ij*_ quantify the coupling of PCP with AB and position, respectively, as described in [20]. These couplings are formulated in in terms of AB vectors **p**_*i*_, PCP vectors **q**_*i*_ and a unit vector 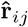 from cell *i* to *j*. The coupling *S*_*ij*_(*AP*) = (**p**_*i*_ × **q**_*i*_) ⋅ (**p**_*j*_ × **q**_*j*_) dynamically maintains the orthogonality of the PCP unit vectors **q**_*i*_ and **q**_*j*_ to their corresponding AB polarity vectors while lateral organization is favored by 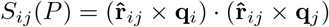. In the absence of any cell shape effects, the coupling between AB and position is given by 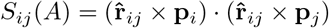, which favours a flat cell sheet. Wedging of cells is introduced into our model by a single deformation parameter *α*, which describes an attractive interaction between the AB polarity unit vectors **p**_*i*_ and **p**_*j*_:

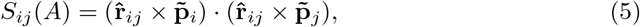

where 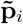 is given by

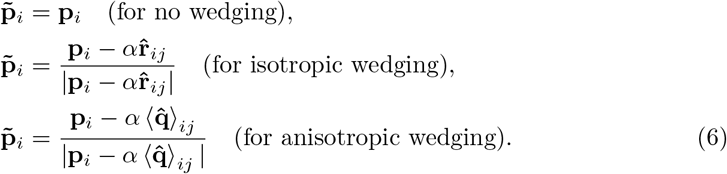

Here, 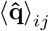 denotes the mean of PCP vectors **q**_*i*_ and **q**_*j*_ belonging to the two interacting cells.

*α* = 0 favors a flat sheet (see Fig 1A–B) whereas a non-zero *α* favours bending of AB polarity vectors towards (or away from) one another and induces curvature in a sheet of cells (Fig 1C–D).

The time development is simulated by overdamped (relaxational) dynamics along the gradient of the above potential, Eq (4). In the case of cell divisions, new daughter cells are placed randomly around the mother cell at a distance of one cell radius.

The source code for the simulations is available on GitHub [39].

### Parameter estimation and robustness

We have tested the robustness of our approach on a number of model cases and find that, for example, *budding* can be reproduced with a broad range of wedging parameters, *α* ∈ [0.1, 0.6] and for diverse PCP coupling strengths *λ*_3_ ∈ [0.8, 0.14]. For these intervals, the budding is qualitatively similar to that illustrated in Fig 2A.

We further explore our model by re-instating dimensions in the formulation of the potential and the equation of motion and estimating dimensionful quantities. With dimensions reinstated, the pair-potential takes the form

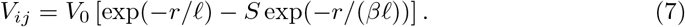

The overdamped equation of motion (without noise) becomes

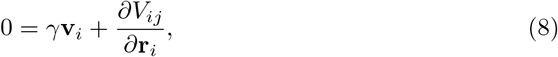

where **v**_*i*_ = *∂***r**_*i*_/*∂t*. We now introduce dimensionless (tilded) parameters

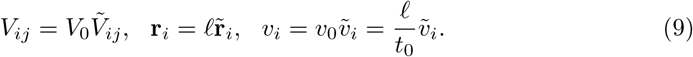

and insert the dimensionless parameters in our equation of motion

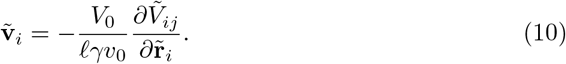

We expect 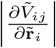, and thus also 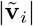 to be of order 1 in sheet-orthogonal extrusion. In [40], a typical value for the dynamical viscosity *µ* was reported to be on the order of 250Pa s. This can be related to the coefficient *γ* by Stokes’ Law of viscous drag, *γ* = 6*πµℓ*. We now compare our model with epithelial cell extrusion and use the typical cell speed reported in [41], *v*_0_ ≈ 1mm h^*−*1^ and use the typical cell size reported in [42], 2*ℓ* = 13µm. With these numbers, our model predicts a typical extrusion energy on the order of

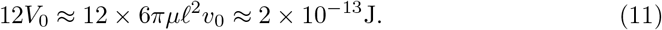

The factor of 12 = 2 × 6 is due to the hexagonal structure of the cell sheet. Note, that our estimate of the extrusion energy is consistent with the finding in [41] for epithelial cell extrusion. Here, an actomyosin ring is measured to exhibit a contraction force of the order of 1kPa, which results in an extrusion energy of the order 1kPa × *ℓ*^3^ ≈ 3 × 10^*−*13^J.

### Modeling neurulation/wrapping

The starting point for our simulation of neurulation is a planar sheet of cells where a line with a width of six cell radii is given non-zero wedging strength |*α*| = *α*_0_ > 0 and all other cells have *α* = 0. The line is centered at *x* = 0 and PCP is initialized orthogonally to this line, along the *x* direction 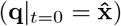. See S4 Fig.

Cell proliferation is simulated as a Poisson process by choosing a rate Γ for *each cell* to divide in each time unit. Only cells at the neuroepithelium-ectoderm interface (defined as cells with *|α| >* 0 who are neighbours of cells with *α* = 0) proliferate (with rate Γ = Γ_0_ *>* 0) while the rest have Γ = 0. Daughter cells inherit all properties of their mother cell and are initiated randomly in a distance of one cell radius from their mother cell.

It should be noted that the initial width of the strip is not particularly important, since wedging will ensure the correct tube width given sufficient proliferation.

All cells in the simulation have the same coupling constants, typically *λ* = (0.6, 0.4, 0). Typical values for Γ_0_ and *α*_0_ are 2.8 × 10^*−*4^ and 0.5. respectively.

### Modeling gastrulation

In our gastrulation simulation, the assignment of PCP and cell wedging is characterized by two radii, describing an annulus (see S5 Fig):

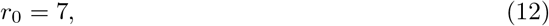

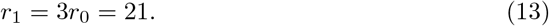

PCP is assigned within the disk Ω_1_ given by

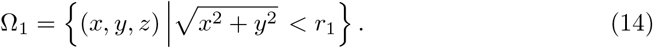

The PCP coupling strength *λ* is taken to be

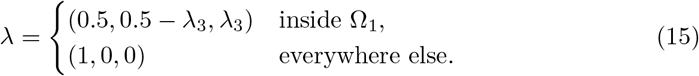

where a typical value for *λ*_3_ is between 0.08 and 0.12.

The PCP vector field **q** is initially assigned so that it spirals around the axis of tube formation (the *z*-axis):

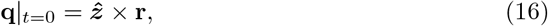

In the gastrulation simulations, the PCP vector field is fixed on a per-cell basis.

Nonzero apical constriction parameter *α* is assigned in an annulus Ω_2_, which shares its outer radius with the disk Ω_1_:

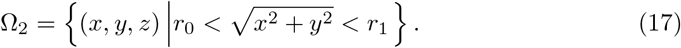

The magnitude of *α* for the cells in Ω_2_ is taken as 0.4:

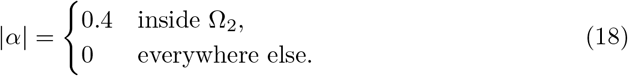

The regions Ω_1_ and Ω_2_ are fixed in space and not on a particle basis. The number of particles in this simulation is *N* = 4000.

### Modeling budding from plane

The budding simulation is, apart from global topology, very similar to the gastrulation simulation.

The relevant length parameters are

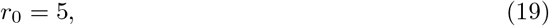

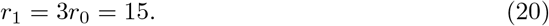

Two regions are correspondingly defined – the disk Ω_1_ and the annulus Ω_2_:

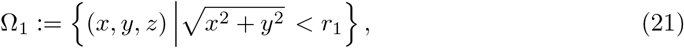

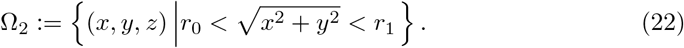

The PCP coupling strength *λ* is taken to be

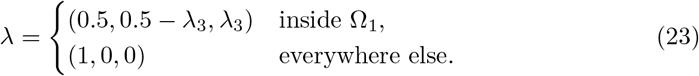

where a typical value for *λ*_3_ is between 0.08 and 0.12.

The PCP vector field **q** is initially assigned so that it spirals around the center of invagination (the origin of coordinates):

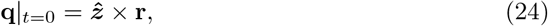

In the gastrulation simulations, the PCP vector field is fixed on a per-cell basis.

Nonzero apical constriction parameter *α* is assigned in the annulus Ω_2_ with magnitude 0.5:

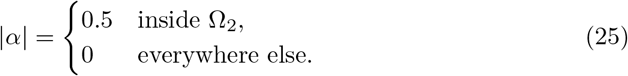

The total number of particles in the simulation is 1384.

## Supporting information

**S1 Fig.**
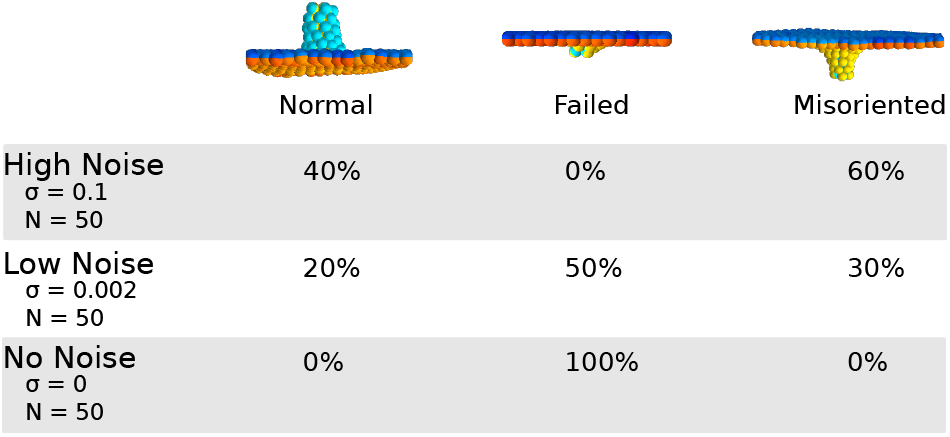
Budding outcomes in the absence of wedging. Budding outcomes without wedging at high and low noise as well as in the absence of noise. The first column shows the proportion of normal initiations of tubulation, the middle column shows failed invaginations while the last column shows misoriented invaginations.

**S2 Fig.**
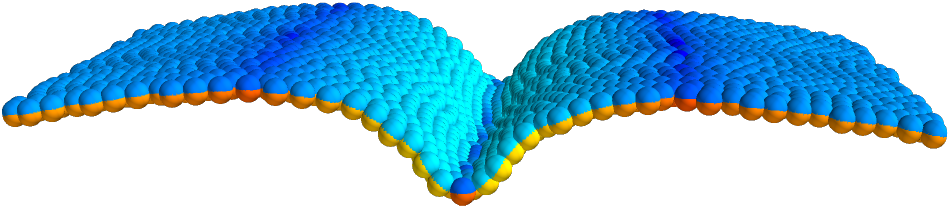
Lack of proliferation. The fate of the neural sheet in our simulations in the absence of proliferation.

**S3 Fig.**
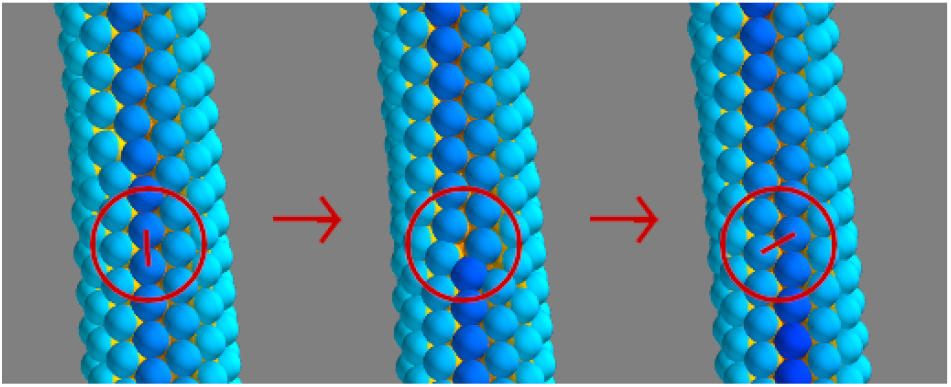
T1 transition induced by wedging.

**S4 Fig.**
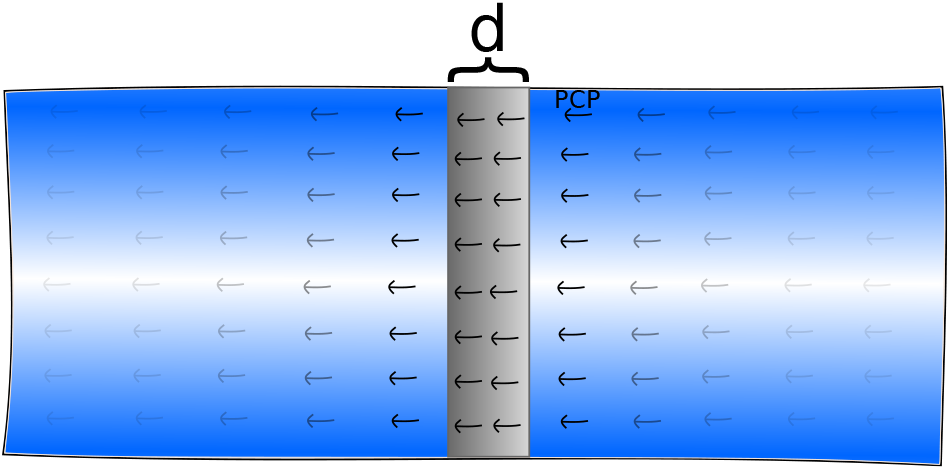
The initial configuration of the cell sheet for neurulation. The initial configurations of the cell sheet for neurulation. Wedging is turned on in a band of width *d* (gray) with PCP running orthogonal to this band.

**S5 Fig.**
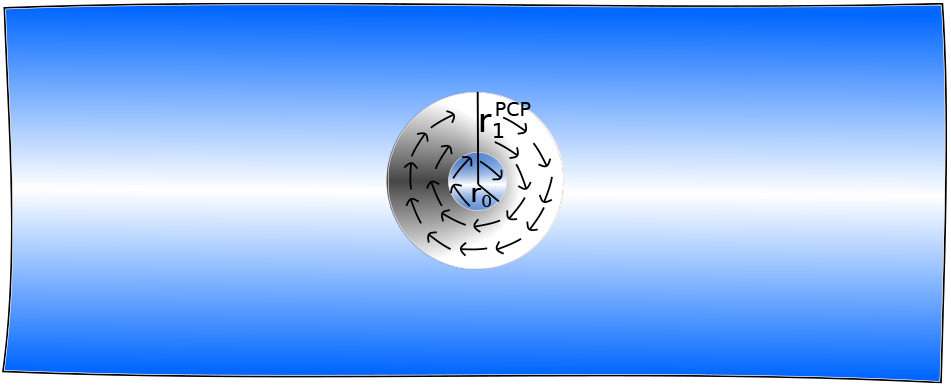
The initial configuration of the cell sheet for budding. The initial configurations of the cell sheet for budding. Wedging is turned on in an annulus (gray) where PCP curls around tangentially.

**S6 Fig.**
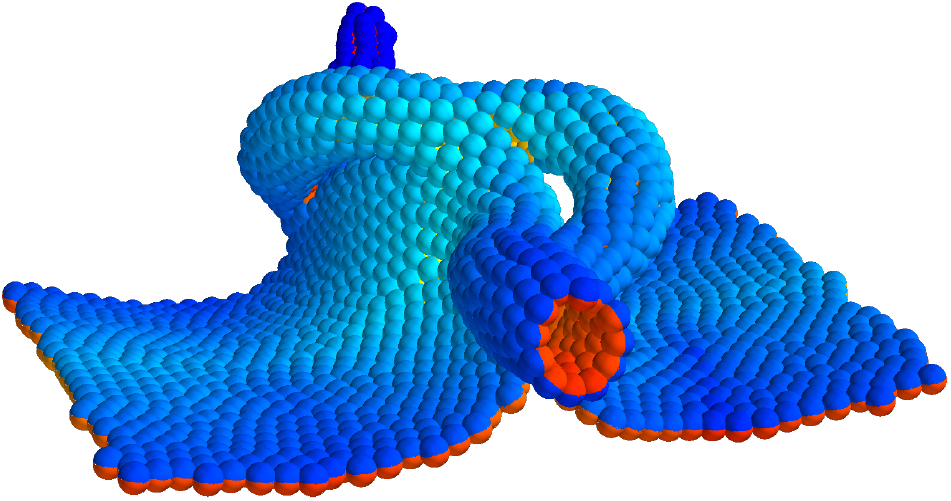
Tube splitting observed with excessive proliferation rate.

S1 Video. Video of budding simulation.

S2 Video. Video of neurulation simulation.

S3 Video. Video of neurulation simulation with isotropic wedging.

S4 Video. Video of a sheet of isotropically apically constricting cells forming a spherical lumen.

S5 Video. Video of sea urchin gastrulation simulation.

## Acknowledgments

This research has received funding from the Danish National Research Foundation (grant number: DNRF116) and the European Research Council under the European Union’s Seventh Framework Programme (FP/2007 2013)/ERC Grant Agreement number 740704.

## References

1. Andrew DJ, Ewald AJ. Morphogenesis of epithelial tubes: Insights into tube formation, elongation, and elaboration. Developmental Biology. 2010;341(1):34–55. doi:https://doi.org/10.1016/j.ydbio.2009.09.024.

2. Sawyer JM, Harrell JR, Shemer G, Sullivan-Brown J, Roh-Johnson M, Goldstein B. Apical constriction: a cell shape change that can drive morphogenesis. Developmental Biology. 2010;341(1):5–19. doi:https://doi.org/10.1016/j.ydbio.2009.09.009.

3. Gutzman JH, Graeden E, Brachmann I, Yamazoe S, Chen JK, Sive H. Basal constriction during midbrain–hindbrain boundary morphogenesis is mediated by Wnt5b and focal adhesion kinase. Biology Open. 2018;7(11):bio034520. doi:https://doi.org/10.1242/bio.034520.

4. Visetsouk MR, Falat EJ, Garde RJ, Wendlick JL, Gutzman JH. Basal epithelial tissue folding is mediated by differential regulation of microtubules. Development. 2018;145(22):dev167031. doi:https://doi.org/10.1242/dev.167031.

5. Chung S, Kim S, Andrew DJ. Uncoupling apical constriction from tissue invagination. eLife. 2017;6:e22235. doi:https://doi.org/10.7554/eLife.22235.

6. Paluch E, Heisenberg CP. Biology and physics of cell shape changes in development. Current Biology. 2009;19(17):R790–R799. doi:https://doi.org/10.1016/j.cub.2009.07.029.

7. Sanchez-Corrales YE, Blanchard GB, Röper K. Radially patterned cell behaviours during tube budding from an epithelium. eLife. 2018;7:e35717. doi:https://doi.org/10.7554/eLife.35717.

8. Nishimura T, Honda H, Takeichi M. Planar cell polarity links axes of spatial dynamics in neural-tube closure. Cell. 2012;149(5):1084–1097. doi:https://doi.org/10.1016/j.cell.2012.04.021.

9. Sweeton D, Parks S, Costa M, Wieschaus E. Gastrulation in Drosophila: the formation of the ventral furrow and posterior midgut invaginations. Development. 1991;112(3):775–789.

10. Martin AC, Gelbart M, Fernandez-Gonzalez R, Kaschube M, Wieschaus EF. Integration of contractile forces during tissue invagination. The Journal of Cell Biology. 2010;188(5):735–749. doi:https://doi.org/10.1083/jcb.200910099.

11. Ossipova O, Kim K, Lake BB, Itoh K, Ioannou A, Sokol SY. Role of Rab11 in planar cell polarity and apical constriction during vertebrate neural tube closure. Nature Communications. 2014;5:3734. doi:https://doi.org/10.1038/ncomms4734.

12. Lee JY, Marston DJ, Walston T, Hardin J, Halberstadt A, Goldstein B. Wnt/Frizzled signaling controls C. elegans gastrulation by activating actomyosin contractility. Current Biology. 2006;16(20):1986–1997. doi:https://doi.org/10.1016/j.cub.2006.08.090.

13. Croce J, Duloquin L, Lhomond G, McClay DR, Gache C. Frizzled5/8 is required in secondary mesenchyme cells to initiate archenteron invagination during sea urchin development. Development. 2006;133(3):547–557. doi:https://doi.org/10.1242/dev.02218.

14. Choi SC, Sokol SY. The involvement of lethal giant larvae and Wnt signaling in bottle cell formation in Xenopus embryos. Developmental Biology. 2009;336(1):68–75. doi:https://doi.org/10.1016/j.ydbio.2009.09.033.

15. Collinet C, Rauzi M, Lenne PF, Lecuit T. Local and tissue-scale forces drive oriented junction growth during tissue extension. Nature Cell Biology. 2015;17(10):1247. doi:https://doi.org/10.1038/ncb3226.

16. Belmonte JM, Swat MH, Glazier JA. Filopodial-tension model of convergent-extension of tissues. PLoS Computational Biology. 2016;12(6):e1004952. doi:https://doi.org/10.1371/journal.pcbi.1004952.

17. Spahn P, Reuter R. A vertex model of Drosophila ventral furrow formation. PLoS One. 2013;8(9):e75051. doi:https://doi.org/10.1371/journal.pone.0075051.

18. Kim HY, Varner VD, Nelson CM. Apical constriction initiates new bud formation during monopodial branching of the embryonic chicken lung. Development. 2013;140(15):3146–3155. doi:https://doi.org/10.1242/dev.093682.

19. Inoue Y, Suzuki M, Watanabe T, Yasue N, Tateo I, Adachi T, et al. Mechanical roles of apical constriction, cell elongation, and cell migration during neural tube formation in Xenopus. Biomechanics and Modeling in Mechanobiology. 2016;15(6):1733–1746. doi:https://doi.org/10.1007/s10237-016-0794-1.

20. Nissen SB, Rønhild S, Trusina A, Sneppen K. Theoretical tool bridging cell polarities with development of robust morphologies. eLife. 2018;7:e38407. doi:https://doi.org/10.7554/eLife.38407.

21. Habib SJ, Chen BC, Tsai FC, Anastassiadis K, Meyer T, Betzig E, et al. A localized Wnt signal orients asymmetric stem cell division in vitro. Science. 2013;339(6126):1445–1448. doi:https://doi.org/10.1126/science.1231077.

22. Loh KM, van Amerongen R, Nusse R. Generating cellular diversity and spatial form: Wnt signaling and the evolution of multicellular animals. Developmental Cell. 2016;38(6):643–655. doi:https://doi.org/10.1016/j.devcel.2016.08.011.

23. Kominami T, Takata H. Gastrulation in the sea urchin embryo: a model system for analyzing the morphogenesis of a monolayered epithelium. Development, Growth & Differentiation. 2004;46(4):309–326. doi:https://doi.org/10.1111/j.1440-169x.2004.00755.x.

24. Kimberly EL, Hardin J. Bottle cells are required for the initiation of primary invagination in the sea urchin embryo. Developmental Biology. 1998;204(1):235–250. doi:https://doi.org/10.1006/dbio.1998.9075.

25. Lyons DC, Kaltenbach SL, McClay DR. Morphogenesis in sea urchin embryos: linking cellular events to gene regulatory network states. Wiley Interdisciplinary Reviews: Developmental Biology. 2012;1(2):231–252. doi:https://doi.org/10.1002/wdev.18.

26. Wallingford JB, Fraser SE, Harland RM. Convergent extension: the molecular control of polarized cell movement during embryonic development. Developmental Cell. 2002;2(6):695–706. doi:https://doi.org/10.1016/S1534-5807(02)00197-1.

27. Galea GL, Nychyk O, Mole MA, Moulding D, Savery D, Nikolopoulou E, et al. Vangl2 disruption alters the biomechanics of late spinal neurulation leading to spina bifida in mouse embryos. Disease models & mechanisms. 2018;11(3):dmm032219. doi:https://doi.org/10.1242/dmm.032219.

28. Smith JL, Schoenwolf G C. Neurulation: coming to closure. Trends in Neurosciences. 1997;20(11):510–517.

29. McShane SG, Molè MA, Savery D, Greene ND, Tam PP, Copp AJ. Cellular basis of neuroepithelial bending during mouse spinal neural tube closure. Developmental Biology. 2015;404(2):113–124. doi:https://doi.org/10.1016/j.ydbio.2015.06.003.

30. Copp AJ, Brook FA, Roberts HJ. A cell-type-specific abnormality of cell proliferation in mutant (curly tail) mouse embryos developing spinal neural tube defects. Development. 1988;104(2):285–295.

31. Tassabehji M, Read AP, Newton VE, Patton M, Gruss P, Harris R, et al. Mutations in the PAX3 gene causing Waardenburg syndrome type 1 and type 2. Nature Genetics. 1993;3(1):26. doi:https://doi.org/10.1038/ng0193-26.

32. Baldwin CT, Lipsky NR, Hoth CF, Cohen T, Mamuya W, Milunsky A. Mutations in PAX3 associated with Waardenburg syndrome type I. Human Mutation. 1994;3(3):205–211. doi:https://doi.org/10.1002/humu.1380030306.

33. Wu TF, Yao YL, Lai IL, Lai CC, Lin PL, Yang WM. Loading of PAX3 to mitotic chromosomes is mediated by arginine methylation and associated with Waardenburg syndrome. Journal of Biological Chemistry. 2015;290(33):20556–20564. doi:https://doi.org/10.1074/jbc.M114.607713.

34. Röper K. Anisotropy of Crumbs and aPKC drives myosin cable assembly during tube formation. Developmental Cell. 2012;23(5):939–953. doi:https://doi.org/10.1016/j.devcel.2012.09.013.

35. Pirraglia C, Walters J, Myat MM. Pak1 control of E-cadherin endocytosis regulates salivary gland lumen size and shape. Development. 2010;137(24):4177–4189. doi:https://doi.org/10.1242/dev.048827.

36. Leptin M, Roth S. Autonomy and non-autonomy in Drosophila mesoderm determination and morphogenesis. Development. 1994;120(4):853–859.

37. Li Y, Muffat J, Omer A, Bosch I, Lancaster MA, Sur M, et al. Induction of expansion and folding in human cerebral organoids. Cell Stem Cell. 2017;20(3):385–396. doi:https://doi.org/10.1016/j.stem.2016.11.017.

38. Kerszberg M, Changeux JP. A simple molecular model of neurulation. BioEssays. 1998;20(9):758–770. doi:https://doi.org/10.1002/(SICI)1521-1878(199809)20:9¡758::AID-BIES9¿3.0.CO;2-C.

39. Nielsen BF. OrganogenesisPCP; 2019. https://github.com/BjarkeFN/OrganogenesisPCP.

40. Eskandari H, Salcudean SE. Characterization of the viscosity and elasticity in soft tissue using dynamic finite elements. In: 2008 30th Annual International Conference of the IEEE Engineering in Medicine and Biology Society. IEEE; 2008. p. 5573–5576.

41. Yamada S, Iino T, Bessho Y, Hosokawa Y, Matsui T. Quantitative analysis of mechanical force required for cell extrusion in zebrafish embryonic epithelia. Biology Open. 2017;6(10):1575–1580. doi:https://doi.org/10.1242/bio.027847.

42. Brown N, Bron AJ. An estimate of the human lens epithelial cell size in vivo. Experimental Eye Research. 1987;44(6):899–906.

